# Which are the most heat-tolerant animals? Insights from a Mediterranean lepismatid under thermal stress in the context of climate change

**DOI:** 10.1101/2025.11.25.690525

**Authors:** Elena Fernández-Vizcaíno, Rafael Molero-Baltanás, José Carbonell, Miquel Gaju-Ricart, Agustín Camacho

## Abstract

Measuring behavioural and physiological thermal limits is crucial to understanding how they interact with the environment under a climate change scenario. We experimentally assessed the effects of acclimation on sequentially measured voluntary (VTmax), critical (CTmax), and upper thermal limit (UTL) limits in the Mediterranean silverfish *Sceletolepisma guadianicum*. Individuals were acclimated for six days at either 25°C (n=32) or 35°C (n=29) and heated at ∼0.5°C min^−1^, and VTmax, CTmax, and lethal limits were recorded. *S. guadianicum* exhibited some of the highest thermal limits reported to date among terrestrial arthropods. VTmax showed limited (1.04 °C) but statistically detectable plasticity, increasing with high acclimation temperature and heating rate, whereas CTmax rate and lethal limits remained unchanged. We provide hypotheses explaining the co-ocurrence of exceptional heat tolerance levels together with their reduced plasticity in this and other extremely heat-tolerant species.

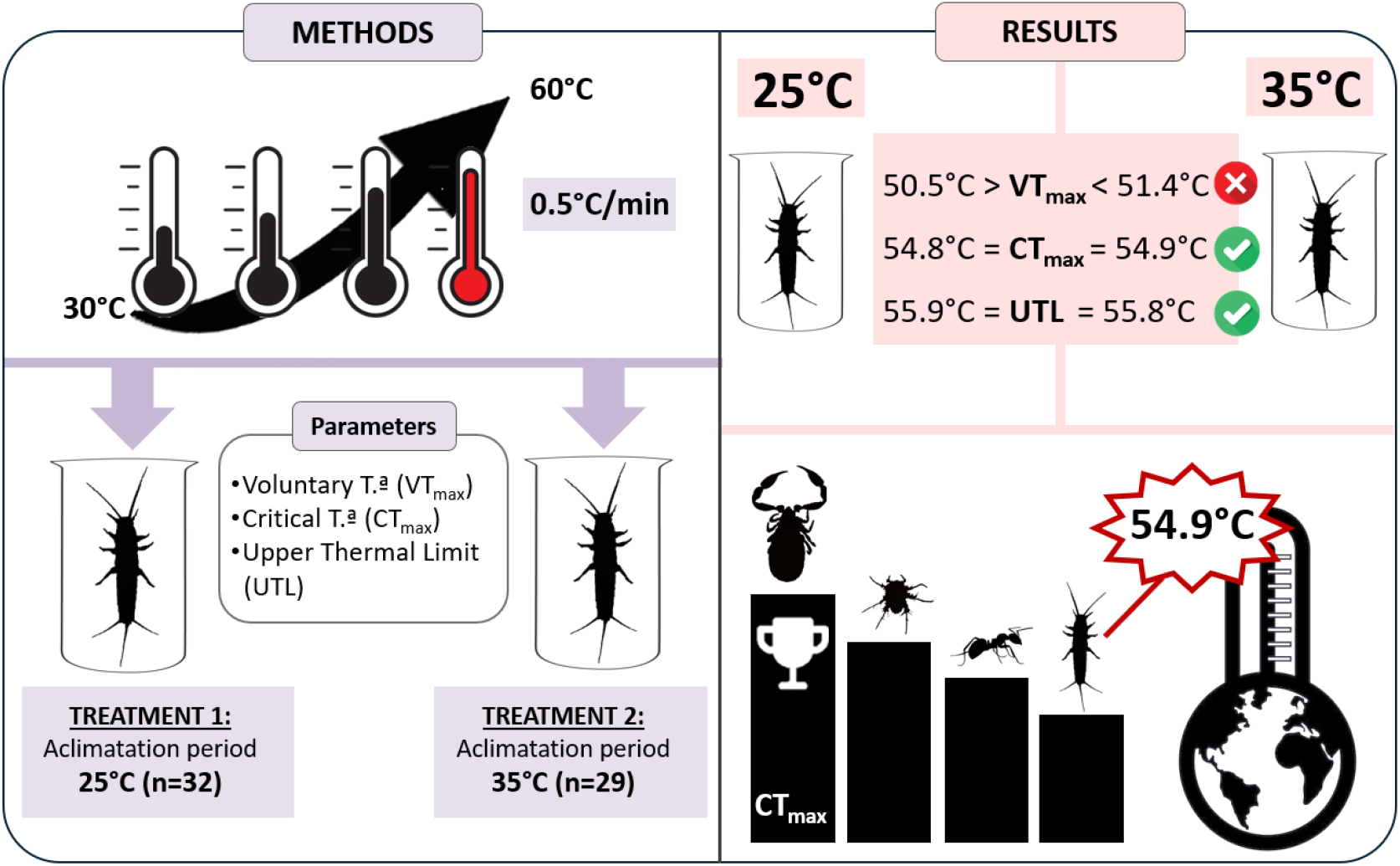

## 1. INTRODUCTION

Extreme heat can disrupt vital cellular functions, reducing performance and jeopardising survival (Angilletta Jr., 2009; Gangloff and Telemeco, 2018). The thermal limits of a species represent its physiological capacity to withstand high temperatures and, depending on environmental variability, can restrict its geographical distribution to sites where animals can defend these limits from climatic conditions (Camacho et al., 2024; Jeffree and Jeffree, 1994). Consequently, extensive interest is directed to establish animal heat-tolerance limits and exploring their interconnections with acclimatory mechanisms and the thermal environments where tolerance manifests‘(Enriquez-Urzelai et al., 2020; Pinsky et al., 2019).

Our knowledge about global animal upper thermal limits derives from a handful of species with exceptional heat tolerance inhabiting extreme environments such as deserts, hot springs, and hydrothermal vents (Bates et al., 2010; Hoffmann et al., 2013; Pikuta et al., 2007). The group Chelicerata harbours the most heat tolerant species, including the mite *Paratarsotomus macropalpis* (Banks, 1916) *with* a CTmax of 60°C (Wu and Wright, 2015) and the pseudoscorpion *Eremogarypus perfectus* Beier, 1962, with a heat stupor point of 65°C (Heurtault and Vannier, 1990). Other hight tolerant insects include the Sahara Desert ant, foraging under near 55°C sun (Myneni et al., 1997), while there are marine animals (E.g., polychaetes, Sipunculida) living near vents with temperatures up to 90°C (Cary et al., 1998; Chevaldonné et al., 2000; Lee, 2003). Likewise, our understanding of how the heat tolerance of these extremophiles responds to variation in the thermal environment, such as acclimation temperature and heating rate, is beginning to gain importance to estimate the implications that natural thermal dynamics may have for animals’ responses (Braz-Mota and Luis Val, 2024; Campos et al., 2019; Sunday et al., 2011). Heat tolerance has traditionally been assessed using the Upper Thermal Limit (UTL), defined as the temperature at which organisms die, and the Critical Thermal Maximum (CTmax), denoting the temperature at which physiological failure leads to loss of motor control in ectotherms (Lutterschmidt and Hutchison, 1997). Nevertheless, many animals avoid lethal conditions behaviourally (Desforges et al., 2023) or are killed by longer exposures to lower temperatures (Rezende and Bacigalupe, 2015). Thus, the measurement of the voluntary thermal maximum (VTmax) has been developed as an alternative measure, indicating the temperature at which organisms actively seek to avoid heat stress (Camacho et al., 2018). Measuring Ctmax and Vtmax sequentially offers a more comprehensive understanding of both physiological limits and behavioural strategies, as exemplified in leaf-cutting ants, where both thresholds showed complex, non-linear relations (Lima et al., 2022).

Besides ecological factors intrinsic to the species and the individual (e.g., body size, sex), thermal limits such as CTmax and VTmax can also vary with environmental (e.g.Bota-Sierra et al., 2022) or experimental conditions, including the initial temperature before heating, heating rate, and acclimation period, reflecting underlying thermoregulatory mechanisms (Castillo-Pérez et al., 2022; Chown et al., 2009; Terblanche et al., 2007). In insects, it has been demonstrated that resistance to short periods of heat stress can adjust depending on prior thermal conditions experienced (Kristensen et al., 2008; Kristensen et al., 2016; Xing and Zhao, 2022). Therefore, brief exposures to non-lethal but elevated temperatures can enhance subsequent heat resistance in terrestrial ectotherms, demonstrating some plasticity in thermal limits (Hoffmann et al., 2013; Weaving et al., 2022).

Lepismatids, a basal insect group, are widely distributed across warm and temperate regions, showing pronounced diversity in arid tropical habitats, suggesting a high heat-tolerance (Molero-Baltanás et al., 2024a; Molero-Baltanás et al., 2024b). However, the heat tolerance of lepismatids, or their acclimatory responses, remains almost unstudied compared to other arthropods (as reviewed in Camacho et al., 2024 and Weaving et al., 2022). This study provides the first experimental assessment of behavioural (VTmax) and physiological (CTmax and ULT) thermal limits in a lepismatid, controlling for heating rate and sex, and examines the influence of acclimation on these limits. Specifically, we studied *Sceletolepisma guadianicum* (Mendes, 1993), which inhabits sun-exposed Mediterranean rocky slopes where surface temperatures often exceed 60 °C (pers. obs.) and remains active and reproductive during warm seasons, suggesting exceptional heat tolerance (Molero-Baltanás et al., 2000; 2015).

## 2. MATERIAL AND METHODS

### 1 Experimental design

Sixty-five *S. guadianicum* adults were collected in June and October 2025 in Alcolea (Córdoba, Spain) and housed under natural shelter conditions (gravel), with water and plant detritus ad libitum. Four individuals were euthanized by freezing and subsequently stored for later use as biothermal probes. The remaining were randomly assigned to two acclimation treatments (25°C, n=32; 35°C, n=29) for six days, with humidity maintained via cotton-plugged water tubes and a 12:12 h light–dark cycle. A setup based on Lima et al. (2022) was used (Fig.1) which consists of a 1⍰L insulated container crossed by five glass tubes (10⍰cm long /1.5cm diameter). The tubes’ central part pass through the water filled contained and are fully submerged under water, while a 3⍰cm long end is exposed to ambient air, outside the container, creating a thermal refuge. A plunger allows controlling the access to the thermal refuge. Water heating was achieved via a submerged insulated resistance cable. Start temperature was 30°C and heating rates varied from 0.4 to 0.7°C/min. Each trial contained four live individuals (two per treatment) plus one dead body with an inserted thermocouple, connected to a PicoLog TC-08 logger, so as to mimic heat exchange of the specimens’ bodies with the heating device, during temperature measurement.

**Figure 1.**
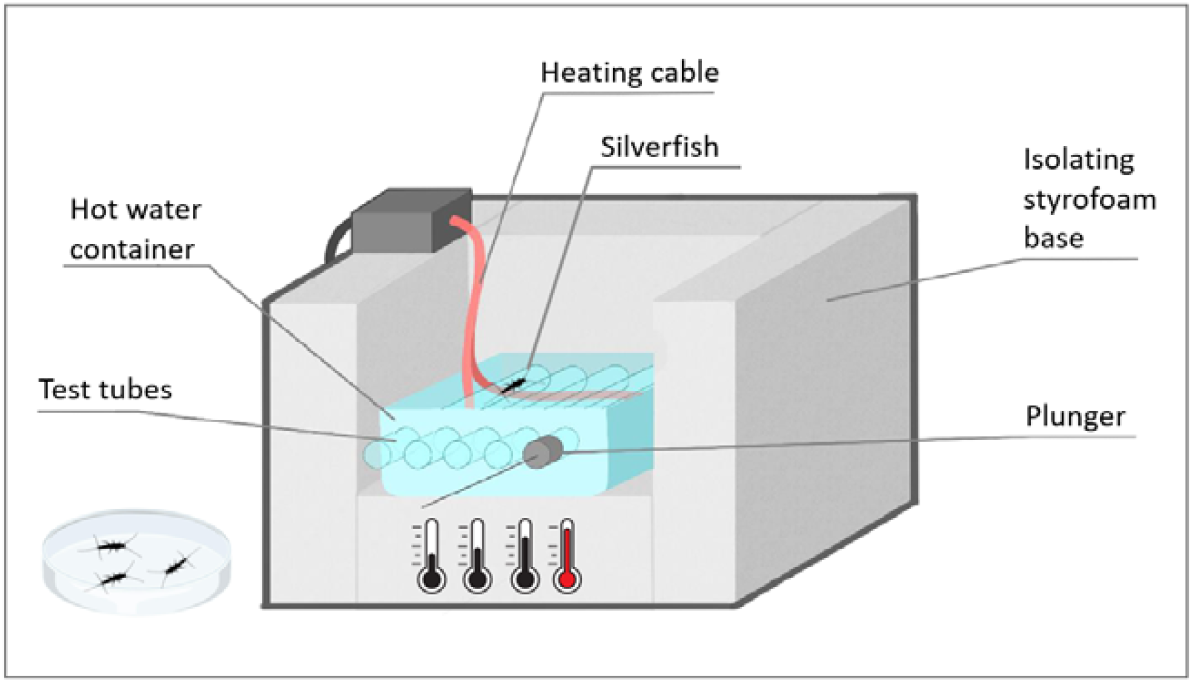
Schematic representation of the thermal tolerance setup.

**Figure 2.**
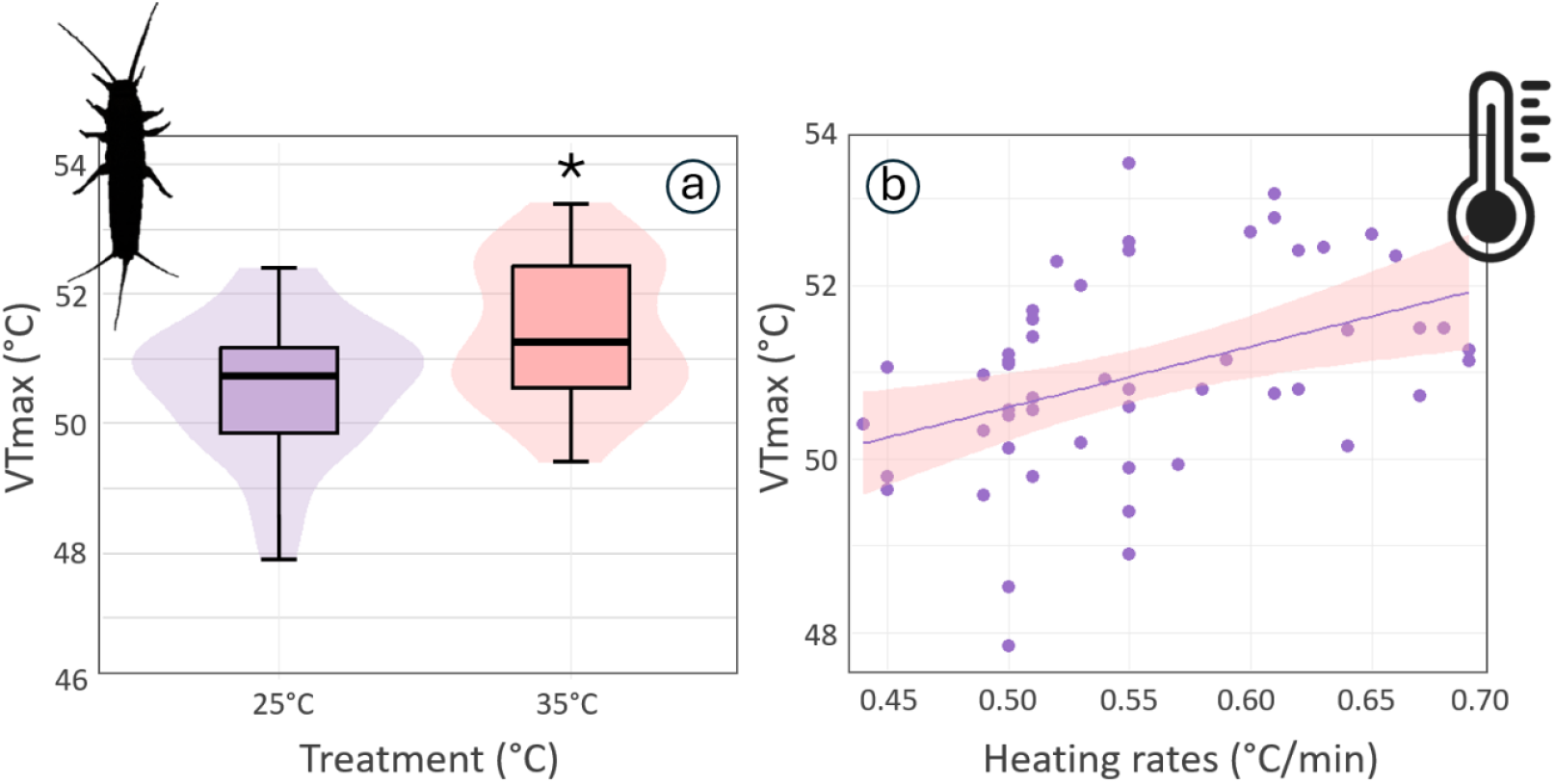
(a) VTmax distribution between acclimation treatments: Treatment 1 at 25°C (purple) and Treatment 2 at 35°C (red). Boxplots show medians and interquartile ranges; violin plots represent the data distribution. Relationship between heating rate and VTmax. Purple points show individual values, the line represents the linear fit, and the pastel red shading indicates the 95% confidence interval.

One silverfish was placed in the submerged section of each glass tube. Individuals remained mostly inactive until the temperature rose, prompting movement towards the thermal refuge. The temperature at which a specimen found the refuge and remained inside it was recorded as the voluntary thermal maximum (VTmax). The plunger was then used to move the specimen, and block access to, the thermal refuge, forcing the animal to remain in the heated zone. As temperature increased, individuals progressively lost coordination and entered a heat-induced coma, characterized by immobility, laterally positioned antennae, and splayed legs. This temperature was recorded as the critical thermal maximum (CTmax). Specimens were then transferred to Petri dishes at 25°C. After 24⍰h, survival was assessed and sex was determined via ovipositor presence. Recovery confirmed CTmax; whereas failure to recover indicated Upper Thermal Limit (UTL; n=16; Lutterschmidt and Hutchison, 1997).

### 2. Statistical analysis

We used R software (R Core Team, 2020) for statistical analyses. The normality of residuals for all numerical response variables was assessed using Shapiro–Wilk tests. Differences between treatment groups were analyzed using analysis of variance (ANOVA), fitting the response variables to models assuming a normal distribution. We fitted separated linear models using analysis of variance (ANOVA), relating each responses in thermal traits (either VTmax, CTmax or UTL) with acclimation treatment, heating rate and sex. For each trait’s model, we performed a model selection procedure, comparing full models with sequentially simpler model, until the null model (i.e., including only intercepts), through a backwards selection procedure. Model comparisons were based on the Akaike Information Criterion (AIC), with lower values indicating better fit, and models differing by at least two AIC units (ΔAIC ≥ 2) considered substantially better supported (Burnham and Anderson, 2004). To interpret the best fitting model, the statistical significance level was set at p < 0.05, considering p-values between 0.05 and 0.1 as marginally significant.

## 3. RESULTS AND DISCUSSION

The mean VTmax ± SD was 50.92 ± 1.16°C, CTmax was 54.45 ± 0.68°C, and lethal threshold temperature was 55.85 ± 0.56°C (Table 1). Model selection indicated that the full model including acclimation treatment, sex, and heating rate best explained variation in both VTmax and CTmax (AIC= 162.18 and 88.36, respectively) whereas the intercept-only model best explained variation in the UTL (AIC= 30.23). Specimens acclimated to higher temperatures (35°C) for six days showed higher VTmax (51.4 ± 1.09°C) than those at 25°C (50.5 ± 1.07°C; F = 11.60, d.f. = 1, p = 0.001). In contrast, CTmax was not affected by treatment or sex, but showed a marginal decrease at higher heating rates (F = 3.83, d.f. = 1, p = 0.058). No effects of treatment, sex, or heating rate were detected for lethal temperature.

**Table 1.** Mean, minimum (Min), maximum (Max), and standard deviation (±SD) values for VTMax, CTMax and UTL per treatment during the acclimation period. Values in bold and with asterisks (*) are those significantly different from controls (p < 0.05).

| | Treatment<br>(Acclimation<br>temperature) | N | Mean ($^\circ\text{C}$) | Min ($^\circ\text{C}$) | Max ($^\circ\text{C}$) | SD ($^\circ\text{C}$) |
| --- | --- | --- | --- | --- | --- | --- |
| VTMax ($^\circ\text{C}$) | $25^\circ\text{C}$ | 30 | 50.5 | 47.8 | 52.4 | 1.07 |
| | $35^\circ\text{C}$ | 25 | **51.4\*** | 49.4 | 53.4 | 1.09 |
| CTMax ($^\circ\text{C}$) | $25^\circ\text{C}$ | 20 | 54.8 | 53.4 | 56.6 | 0.78 |
| | $35^\circ\text{C}$ | 21 | 54.9 | 53.8 | 55.7 | 0.57 |
| UTL ($^\circ\text{C}$) | $25^\circ\text{C}$ | 9 | 55.9 | 55.1 | 56.7 | 0.55 |
| | $35^\circ\text{C}$ | 5 | 55.8 | 55.0 | 56.7 | 0.63 |

The highest known heat tolerances among terrestrial animals have been recorded in chelicerates, such as the pseudoscorpion *E. perfectus* (heat stupor point = 65°C; Heurtault & Vannier, 1990) and the mite *P. macropalpis* (CTmax = 60°C; Wu & Wright, 2015). Among insects, the Australian desert ant *Melophorus bagoti* Lubbock, 1883 (CTmax = 56.7°C; Christian & Morton, 1992), capable of surviving for one hour at 54.0°C, and the Saharan ant *Cataglyphis bicolor* (Fabricius, 1793) (CTmax = 55.1°C; Gehring and Wehner, 1995) show similarly extreme heat resistance. By detecting a CTmax of 54.9°C, this study places the temperate species *S. guadianicum* closely behind these desert specialists, placing it among the most heat-tolerant arthropods ever reported. Comparable values have also been documented in others thermophilic ants, such as *Ocymyrmex velox* Santschi, 1932 from the Namib Desert (54.1°C; Wehner and Wehner, 2011) and the Saharan silver ant *Cataglyphis bombycina* Roger, 1859 (53.6°C; Gehring & Wehner, 1995) and in some dung beetles (53°C Gotcha et al., 2021). Other studies on lepismatids have reported lower thermal limits, such as *Ctenolepisma ciliatum* (Dufour, 1831) (CTmax = 42.28°C), *Ctenolepisma calvum* Ritter, 1910 (CTmax = 41.79°C), and *Lepisma saccharinum* Linnaeus, 1758 (CTmax = 41.79°C;(Perea, 2023)). In both these studies and ours, specimens were heated within a narrow chamber surrounded by water, a method that homogenizes the thermal environment experienced by individuals and prevents the overestimation of heat tolerance typically observed in assays using dry hot plates (Lima et al., 2022). Additional studies on domestic taxa of lepismatids have mainly focused on thermal preferences and growth rather than critical limits. Although they use different methodology, these studies provide useful comparative context. For *Ctenolepisma longicaudata* Brauns 1905, the optimal temperature is reported 24°C (Aak et al., 2019; Heeg, 1967), with mortality occurring within 2 hours 40-44 °C and quickly above 50 °C (Lindsay, 1940). Moreover, Sweetman (1938, 1939) reported that *Thermobia domestica* (Packard, 1873) develops optimally at 37°C, ceases reproduction above 41°C and rarely survives brief exposures near 50°C. In contrast, *Lepisma saccharina* cannot withstand temperatures above 36°C despite inhabiting regions that frequently exceed this threshold (Molero-Baltanás et al., 2014; Sweetman, 1938). Taken together, our results and previous studies (Sweetman, 1938, 1939) support the idea that species inhabiting natural environments can evolve higher thermal tolerance than their domestic relatives.

The exceptional heat tolerance of *S. guadianicum* and its contrast with other thermophilic lepismatids may reflect both evolutionary history and biogeographic background. The genus *Sceletolepisma* Wygodzinsky, 1955 has a broad distribution across southern Africa, the Palearctic region of Asia, and southern Europe, and it includes some of the most thermophilic species known (Molero-Baltanás et al., 2024a). Interestingly, *Sceletolepisma* was recently separated from *Ctenolepisma* (Molero-Baltanás et al., 2024b) and this taxonomic distinction, together with differences from *Lepisma* and *Thermobia*, may help explain the variation in thermal limits (Sweetman, 1938, 1939). These species are typically more nocturnal and occupy buffered microhabitats, which is consistent with the idea that nocturnal species rely on more stable environments, exhibit reduced heat acclimation, and are restricted to resources that remain accessible under such moderated thermal conditions (Esch et al., 2017; Wehner and Wehner, 2011).

Our acclimation treatment produced low but contrasting effects on VTmax, CTmax, and UTL. Individuals acclimated to 35°C exhibited statistically higher VTmax values than those acclimated to 25°C, indicating a modest degree of plasticity in behavioural upper thermal tolerance. The increase in VTmax was approximately one degree, with substantial overlap in tolerance values between treatments. In contrast, unlike previous reports for insect critical limits (e.g. van Heerwaarden et al., 2016; Xing & Zhao, 2022), CTmax exhibited no statistically significant acclimation response, although mean values were slightly higher (≈0.1°C) in the 35°C treatment. Such low acclimatory changes in different heat tolerance parameters in our study is consistent with findings in other insects which inhabit thermally stressful environments year-round (e.g. Aragon-Traverso et al., 2023). VTmax acclimation data are still scarce across both ectotherms and endotherms. However, species from cooler habitats often show larger degrees of plasticity in CTmax (e.g. van Heerwaarden et al., 2016; Xing and Zhao, 2022). These facts support the hypothesis that more stressful habitats leads to the evolution of constitutive heat tolerance traits, concomitantly with weak acclimatory processes that allow the establishment of predictive relationships between heat tolerance and geographic thermal limits in arthropods and other animals (Camacho et al., 2024). Alternatively, Weaving et al. (2022) comprehensive meta-analysis on insect upper thermal limits proposes that plasticity in insect upper thermal limits is generally weak (average change of 0.09°C in thermal limits per 1°C of acclimation), which also agrees with our observations.

Methodological factors such as heating rate are known also to influence thermal limits in arthropods (Castillo-Pérez et al., 2022; Lima et al., 2022; Terblanche et al., 2007). In our study, we found that VTmax increase under faster temperature ramps, agreeing with findings in other insects, such as leaf-cutting ants, where faster heating rates increased both VTmax and CTmax (Lima et al., 2022), and *Drosophila melanogaster* <u>Meigen</u>, 1830, in which slower ramps produced lower CTmax estimates (Terblanche et al., 2007). Behavioural responses likely depend on how insects perceive the risk of overheating, for instance, faster heating rates could prompt an organism to seek thermal refuge at lower temperatures (a lower VTmax) in anticipation of a higher risk of exceeding their CTmax. However, CTmax showed a slight opposite trend to VTmax, decreasing under faster heating rates. Uncorrelated changes in VTmax and CTmax are rare but documented in both intraspecific (Camacho et al., 2018; Lima et al., 2022) and interspecific contexts (Camacho et al., 2023). For example, in the cricket *Gryllus assimilis* (Fabricius, 1775), heating rate modulated VTmax differently between sexes (Díaz-Ricaurte et al., 2022), while Allen et al. (2012)reported opposite effects of heating rate across species: in *Tenebrio molitor* Linnaeus, 1758, faster rates reduced CTmax, whereas in *Cyrtobagous salviniae* Calder and Sands, 1985 they increased it. These different outcomes indicate that thermal limits can change dynamically in response to multiple interacting factors and might derive from yet undetermined specific habitat requirements.

In conclusion, the exceptional heat tolerance of *S. guadianicum* highlights its remarkable resilience to extreme temperatures and challenges current assumptions about the upper thermal limits of Lepismatidae. This species endures temperatures well above those recorded in the field in Córdoba, where nearly 45 °C were reached in the summer of 2025 (AEMET, 2025), indicating a broad physiological safety margin compared with most ectotherms (Sunday et al., 2012). Documenting such overlooked cases is essential for refining our understanding of insect thermal adaptation and improving predictions of species’ responses to climate warming, particularly in Mediterranean ecosystems where extreme events are expected to intensify (Hidalgo-Triana et al., 2023). Additionally, the influence of acclimation temperature and heating rate on upper thermal limits highlights the importance of such studies for assessing insect responses under ongoing climate change.

## 4. COMPETING INTERESTS

The authors declare no competing interests.

## 5. AUTHOR CONTRIBUTIONS

**Elena Fernández-Vizcaíno:** Data curation, Formal analysis, Investigation, Methodology, Visualization, Writing – original draft; **Rafael Molero:** Data curation, Project administration, Methodology, Resources, Writing – review & editing; **José Carbonell:** Conceptualization, Methodology, Investigation, Writing – review & editing; **Miquel Gaju-Ricart:** Resources, Project administration, Methodology, Writing – review & editing; **Agustín Camacho:** Conceptualization, Investigation, Formal analysis, Methodology, Project administration, Writing – review & editing.

## 6. FUNDING

This work is part of the project “*Productivity and biodiversity maintenance in dehesas and woody crops*” (ref. BIOD22_00033_17_PPCB), funded by the Spanish Ministry of Science, Innovation and Universities and the Regional Government of Andalusia.

## 7. ACKNOWLEDGEMENTS

We are thankful to Alise Jandard for her help in developing the first experimental trials to measure the VTmax of lepismatids and to Ana Cárdenas for her support during the initial stages of this work.

## Notes

### Competing Interest Statement

The authors have declared no competing interest.

